# Breast Cancer Clustering Integrating Complete Gene Expression Profiles and Genetic Ancestry

**DOI:** 10.1101/2025.06.05.658154

**Authors:** Johanna Stepanian, Alejandro Mejia-Garcia, Carlos Orozco, Jorge Duitama

## Abstract

Breast cancer (BC) remains the leading cause of cancer-related mortality among women globally. Precise subtyping of BC is critical for optimizing treatment strategies. This study explored the capacity of bulk RNA-seq data to improve breast cancer characterization by analysis of complete expression profiles. We analyzed RNA-seq 274 tumor samples and six healthy tissue samples from diverse geographical origins. Using over 9,800 SNPs directly genotyped from RNA-seq data, we successfully predicted broad genetic ancestry, identifying European, African, Asian, South Asian, and Admixed American origins. Molecular subtyping through PAM50 showed ambiguous classifications for about half of the samples, underscoring the limitations of current molecular diagnostic tools. Unsupervised clustering separated tumors in three main clusters. Cluster C1 demonstrated immune activation and inflammatory response pathways, while C2 highlighted metabolic and immune interaction processes. Cluster C3 exhibited enriched metabolic regulation and adipokine signaling pathways. *In silico* drug sensitivity analysis identified potential therapeutic strategies, including vinorelbine and AZD6482, with cluster-specific efficacy. Our findings emphasize the integration of ancestry-informed data and complete transcriptomic profiles to redefine BC subtyping. These insights offer a foundation for more equitable, ancestry-informed therapeutic strategies and highlight the importance of diversity in cancer research.

## Introduction

Breast cancer (BC) is the first leading cause of death in women around the world (Sung et al., 2021). Subtypes are important to determine the optimal treatment plan for patients (Szymiczek *et al*., 2020). Classic subtypes are determined by the activity of the estrogen receptor (ER), the progesterone receptor (PR), and the human epidermal growth factor 2 (HER2), measured by immunohistochemistry (IHC) (Fragomeni et al., 2018). Based on these biomarkers, the most aggressive subtype is the triple-negative BC (TNBC), which is negative for ER, PR, and HER2 (Joyce et al., 2017; Ye et al., 2016). Cytotoxic chemotherapy is the main effective therapeutic modality for this subtype. However, most patients present side effects including infertility, osteopenia, and heart damage and some patients develop resistance to the treatment (Nedeljković & Damjanović, 2019).

Despite the availability of high throughput gene expression measurements such as RNA-seq, subtyping is currently performed using the IHC classification method, based on microarray data. This method defines five intrinsic subtypes: Luminal A, Luminal B, Normal-like, Basal and HER2 (Perou et al., 2000). Luminal A shows a good prognosis, a low relapse rate, higher survival time, and sensitivity to endocrine therapy (Arpino et al., 2013; Ciriello et al., 2013; Eroles et al., 2012; Haque et al., 2012). It is usually associated with somatic mutations in *PIK3CA, GATA3*, and *MAP3K1* genes, and with overexpression of the cyclin D1 gene (Norum et al., 2014). Luminal B presents a lower sensibility to endocrine treatment and a higher sensitivity to chemotherapy, compared to Luminal A (Goldhirsch et al., 2011; Ignatiadis et al., 2012). It also shows the worst prognosis within the Luminal subtypes (Ades et al., 2014). Many patients with germline mutations in *BRCA1* develop basal tumors (Foulkes et al., 2003; Jung et al., 2016; Mavaddat et al., 2012). These tumors are highly diverse in terms of epidemiological, phenotypic, and molecular characteristics, with different patterns in terms of relapse (Bertucci et al., 2012; Cadoo et al., 2013). Enriched-HER2 tumors show high expression of genes associated with cellular proliferation (Eroles et al., 2012). They also show intermediate expression in luminal genes (*ESR1* y *PGR)*, and low expression of basal genes and proteins (Prat et al., 2015). These tumors have a good response to monoclonal antibody therapy with Trastuzumab, decreasing the death rate in early metastatic states (Eroles et al., 2012). Resistance to the treatment has been related to overexpression in CXCR4 and the loss of PTEN (Dai et al., 2015; Tekesin et al., 2019). Finally, normal-like tumors have a different expression pattern and the worst prognosis for the patient (Dai et al., 2015).

It is known that the most aggressive intrinsic subtypes are more frequent in Latinas, Native American, and African American women compared to European descent women (Zavala et al., 2021). Although specific factors explaining a higher incidence of HER2+ tumors in Latinas are unknown, a positive correlation between the proportion of native american ancestry and HER2 status has been reported (Marker et al., 2020). Higher rates of HER2 gene expression in the HER2+ subtype have also been reported in southeast Asian patients (Su et al., 2011; Li et al, 2019; Parise et al., 2016; Telli et al., 2011). Additionally, in the United States, these ethnic groups have limited access to health services due to multiple cultural and language barriers (Wisniewski & Walker, 2020). While socioeconomic factors contribute to population-based differences in mortality, they do not explain differences in all populations (Roelands *et al*., 2021).

While immunohistochemistry (IHC) remains widely used due to its cost-effectiveness, it fails to fully capture tumor heterogeneity (Choi et al., 2012). Hence, different commercial kits based on gene expression for BC subtyping were developed and are used in clinical practice. One of the most popular methods, known as PAM50, performs supervised clustering of the expression data obtained from a microarray of 50 genes (Ochoa *et al*. 2020, Mathews *et al*., 2019, Tibshirani, *et al*., 2002; Parker *et al*., 2009). Other panels have been designed such as the Oncotype DX assay, which based on the expression patterns of 21 genes classifies patients into three groups based on the recurrence risk: high, intermediate, and low (Ross *et al*., 2008 y Harris *et al*, 2007, Schaafsma *et al*., 2021). A main limitation of the current classification models is that they have been trained using data primarily from European populations. Hence, they overlook the heterogeneity of breast cancer in patients with diverse genetic ancestries (Shah et al., 2017; Troester et al., 2018; Roelands et al., 2021). Differences in tumor biology and subtype distribution across African, Latin American, and Asian populations have been reported, emphasizing the need for a more inclusive approach to molecular classification (Huo et al., 2017; Roelands et al., 2021; Fejerman et al., 2008; Wisniewski & Walker, 2020; Jack et al., 2014; Januszewski et al., 2014; Howlader et al., 2014). Novel methods including a broader set of genes, more balanced training databases, and ancestry-informed markers are needed to improve diagnostic precision and to achieve equitable, personalized treatment strategies for all breast cancer patients.

In this study, we aim to improve breast cancer subtyping by considering ancestry and by using the nearly complete gene expression profiles that can be reconstructed from RNA-seq data. First, we validated that accurate ancestry predictions can be obtained from direct analysis of RNA-seq data. The clustering of 274 human BC tumors with available RNA-seq data provides new groups that can be related to biological functions and responses to drug treatments.

## Results

### Genetic ancestry prediction for breast cancer samples from RNA-seq data

We analyzed RNA-seq data from tumors of 274 breast cancer patients and six healthy tissue controls, retrieved from 36 publicly available studies. Most of these studies were conducted in the United States and Spain (Figure 1). Reads from all samples included in this study were aligned with a mapping rate exceeding 70% to the reference genome (Supplementary Table 1). To assess whether the genetic ancestry of each sample could be evaluated from RNA-seq data alone, we genotyped known variants in coding regions from the 1,000 Genomes Project, which included 1,600 reference female individuals in their project. This resulted in 9,578 single nucleotide variants (SNVs) genotyped in a total of 1,808 individuals with a missing data rate of 1.85% attributed to RNA-seq differences in expression, ranging between 4524 and 9578 genotype calls per sample (Supplementary Table 1).

**Figure 1.**
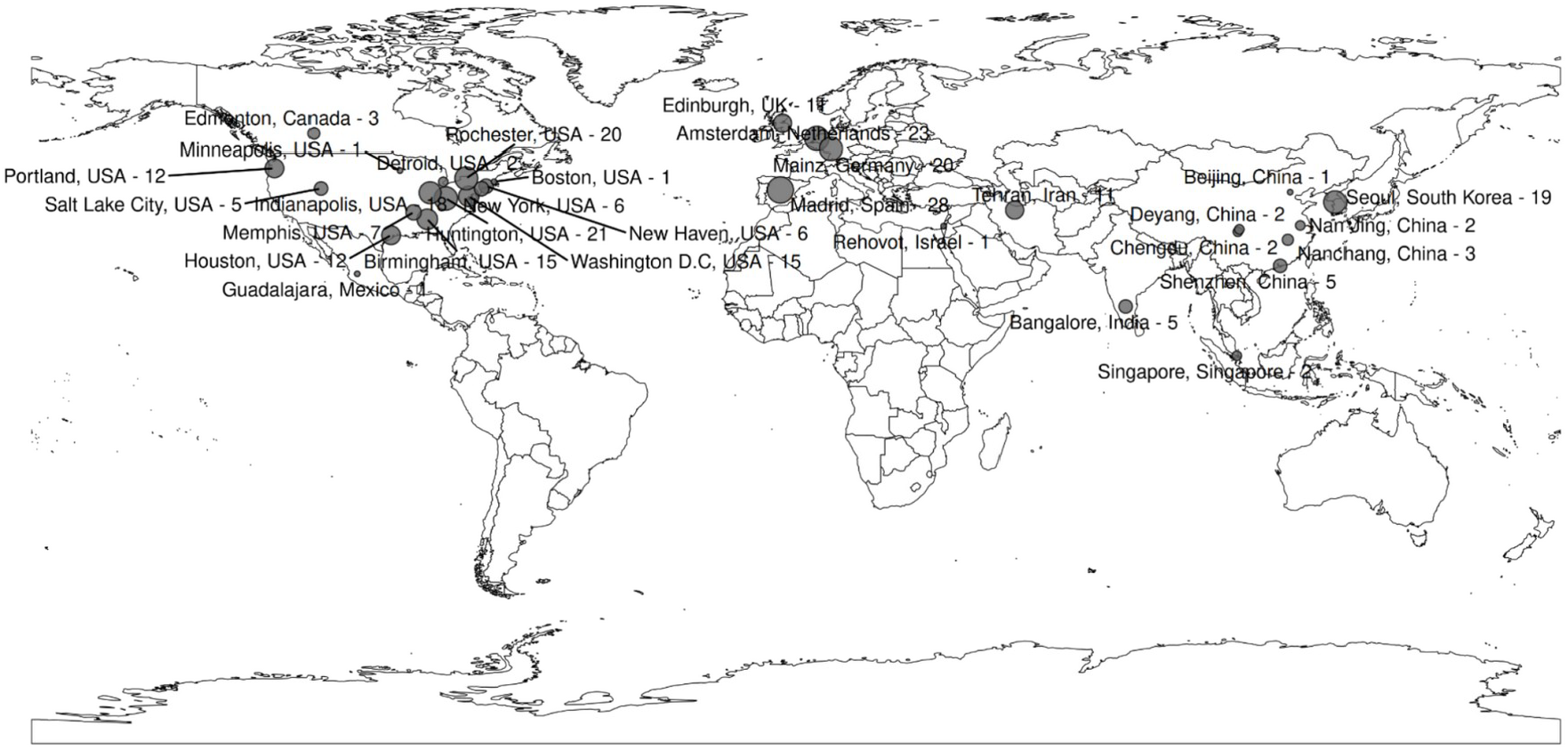
Sample origin from breast cancer women included in the study.

To infer the genetic population of origin of each sample, we performed a maximum likelihood estimation of individual ancestries using the previously called SNVs varying the number of clusters (k parameter) between 1 and 10, using the genomic variation database constructed from RNA-seq data as input. Ancestry was successfully predicted for samples from the 1,000 Genomes Project (Figure 2A). In particular, the African population was split in k=2 given its higher diversity compared to other populations. Asian and European groups split in k=4, and the probably Native American component (AMR, orange) appears in Admixed Americans in k=5, separating them from the EAS population. We chose k=5 to determine the ancestry of the breast cancer samples, considering the previous annotations on human populations. Most of the samples (n=204) were classified as European ancestry (EUR), followed by East Asian (EAS) (n=40), African (AFR) (n=24), South Asian (SAS) (n=7), and Admixed American (AMX) (n = 5) (Figure 2A), which is consistent with the geographic origin of the samples. Individuals from the USA, Israel, Canada, Germany, Spain, the UK, and the Netherlands showed predominantly European ancestry, whereas individuals of Asian origin, including samples from China and South Korea, were classified as EAS. Individuals from Singapore and India clustered predominantly within the SAS population (Figure 2B). Genetic admixture was predicted for individuals from New York, individuals of African origin living in Birmingham, and individuals of Mexican origin (Figure 2C).

**Figure 2.**
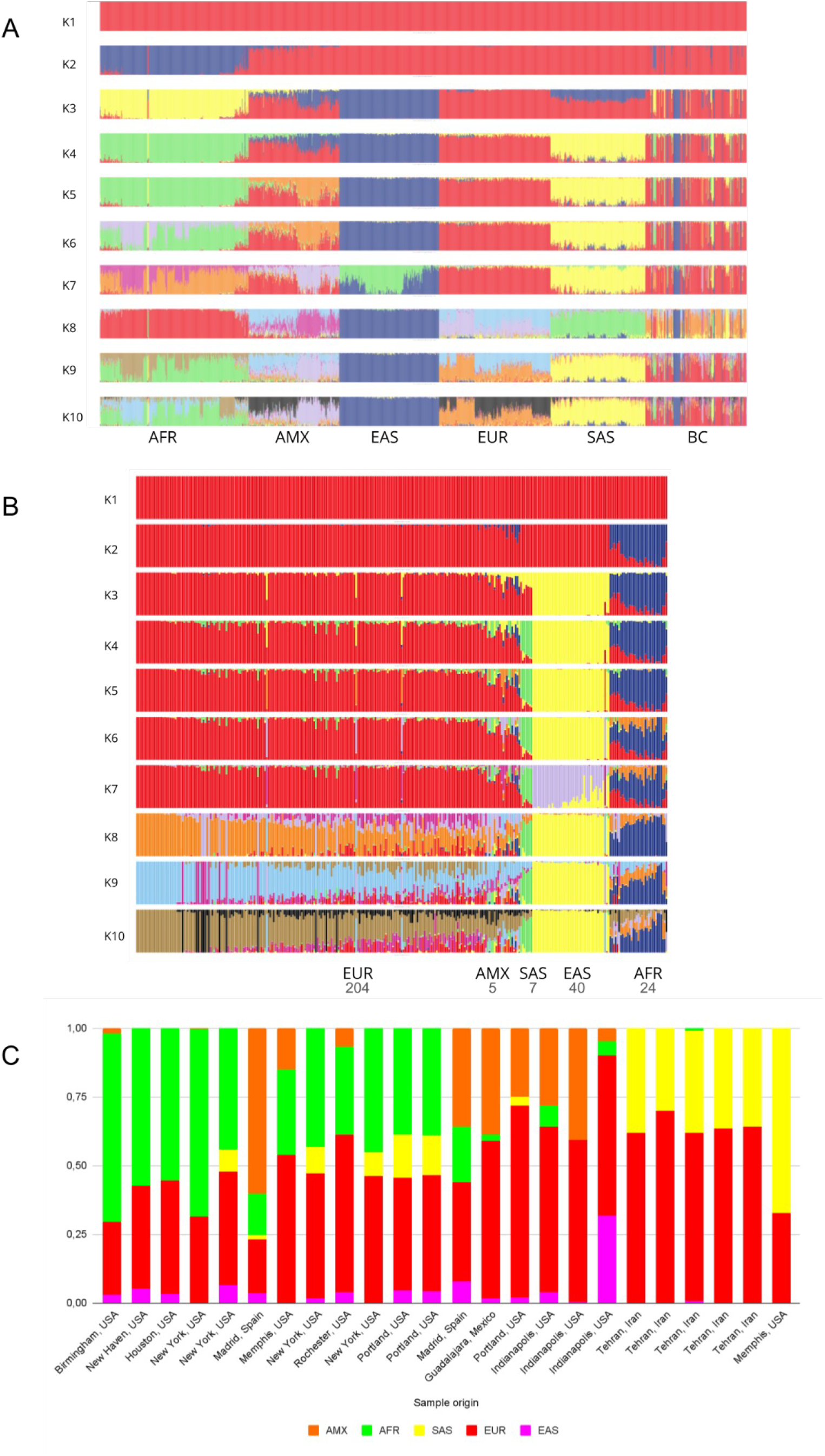
A. ADMIXTURE clustering of individuals from the 1,000 genomes project and breast cancer RNA-seq data. African (AFR), Admixed American (AMX), East Asian (EAS), European (EUR), and South Asian (SAS). B. Ancestry frequency for 280 breast tissue samples by origin. C. Ancestry prediction for the admixed individuals, each column represents an individual.

Besides the five individuals classified as AMX, we observed 24 individuals with admixture patterns. Most of these individuals were originally classified as European. However, they had a membership probability lower than 0.6 to a single population so we re-classified them as: European–African (25%, 6/24), followed by European–South Asian, and European–Admixed Americans with an equal proportion of 21% each (5/24) and European–East Asian (4%, 1/24). Additionally, we found African– Europeans 21% (5/24), South Asian – European (4%, 1/24), Admixed American–African (4%, 1/24).

### Uncertainty in breast cancer subtyping based on the PAM50 panel

We obtained a prediction of breast cancer subtypes for the 274 patients for which we used RNA-seq data using the PAM50 algorithm. If predictions are made based on maximum probability, the most common subtypes were Basal (103 samples) and Luminal A (74 samples); while the other subtypes had a smaller number of samples with an average of 33 samples per subtype. However, if the subtypes are predicted based on a probability greater than 0.6 (Figure 3A), 36% of the samples can not be classified as showing mixed memberships in two or more subtypes (Figure 3B). In particular, 44.59% of the samples classified as Luminal A have a probability higher than 40% of being classified as a Normal-like subtype. A similar situation can be observed for 22.33% of the samples classified as Basal, but having a high probability of being enriched HER2. Considering that for the basal subtype of breast cancer, there are no clear treatment options, about 37.5% of the patients related to the samples included in this study would not have a clear protocol after performing a PAM50 analysis.

**Figure 3.**
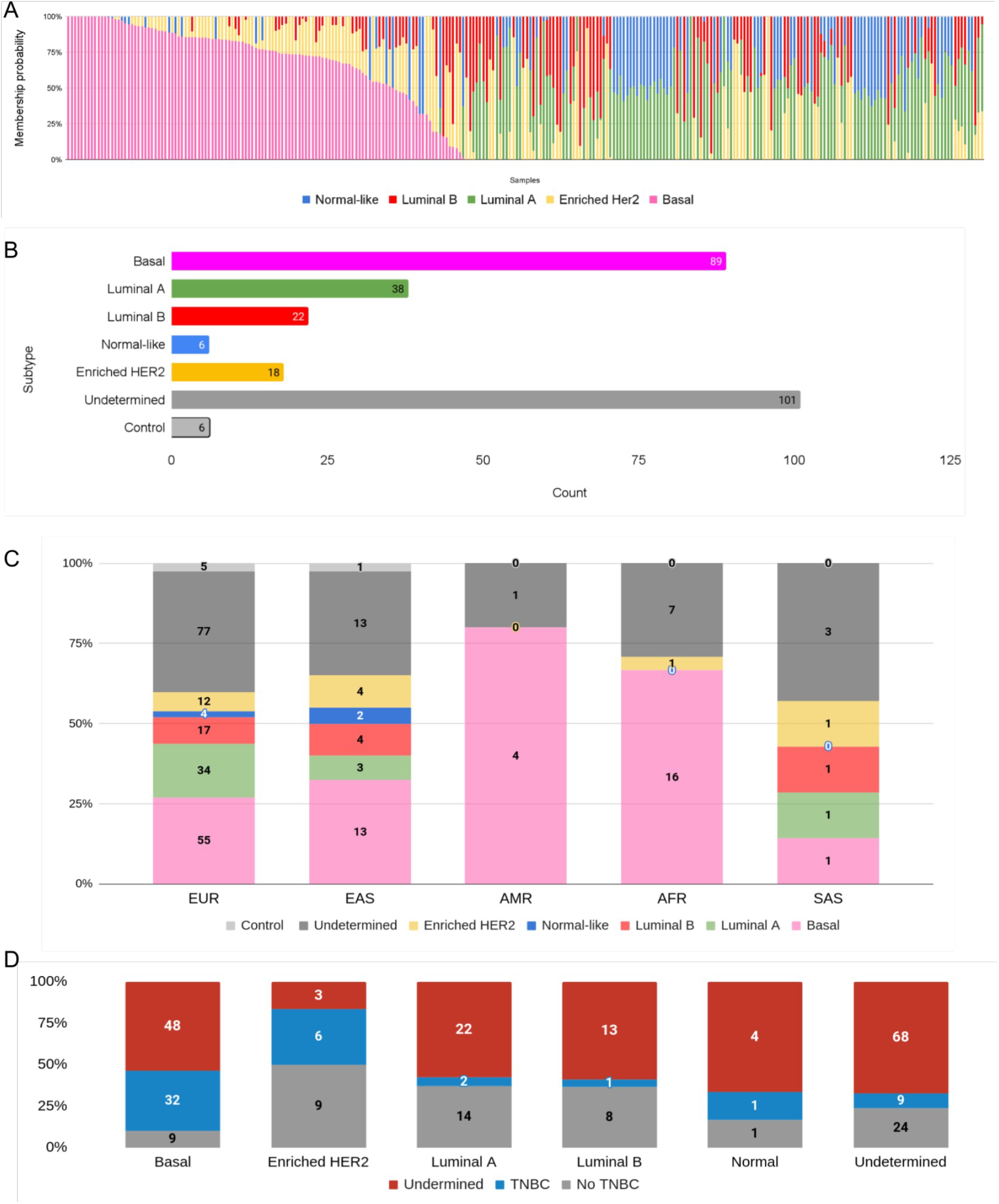
A. Subtype frequency obtained with the PAM50 algorithm for 274 RNA-seq breast cancer tumor and six healthy tissue samples. (LumA = Luminal A, LumB = Luminal B). B. Probability of subtypes for each sample predicted by PAM50. C. Percentage of samples with different subtype predictions discriminated by genetic population. Numbers over the bars show the number of samples. D. PAM50 subtype prediction and its correlation with TNBC status annotated by sample submitter.

We investigated the relationship between the subtypes predicted using the PAM50 algorithm to the genetic populations inferred by ADMIXTURE. In samples from EAS, AMX and AFR the most common subtype is Basal, while in EUR and SAS, a predominant subtype could not be determined due to the large number of samples with undetermined subtypes (Figure 3C). Regarding the correlation between the intrinsic subtype predicted by PAM50 and TNBC status (Figure 3D), although more than half of the samples did not have TNBC as reported metadata, we could assess that most individuals cataloged as TNBC had Basal as predicted subtype (p-value 1.17×10 ^−6^ for a fisher exact test). We finally predicted subtype over control samples, 83% (5 out of 6) were classified as normal-like with at least 30% membership in other subtypes. The remaining control sample was classified as Luminal A.

### Clustering of breast cancer tumors based on complete expression profiles

We built a gene count matrix for the 280 samples (274 breast cancer samples and 6 healthy tissue controls) and constructed a metadata matrix using as features: intrinsic subtype, TNBC status, prediction of genetic population (at k=5) with and without considering admixed individuals, and cancer status. A principal component analysis was performed to evaluate variance and batch effects. Figure 4 shows that the samples seem to cluster in three to four different groups, according to the two first principal components, which take into account 44% of the variance. This clustering does not seem to be related to TNBC status (Figure 4A), or predicted subtype by PAM50 (Figure 4B). Regarding genetic ancestry, while EUR individuals were spread out through all the plots, all SAS samples showed a particular pattern of clustering in the upper right cluster (Figure 4C). Admixed samples (filled circles, Figure D) were present in two clusters (Figure 4D).

**Figure 4.**
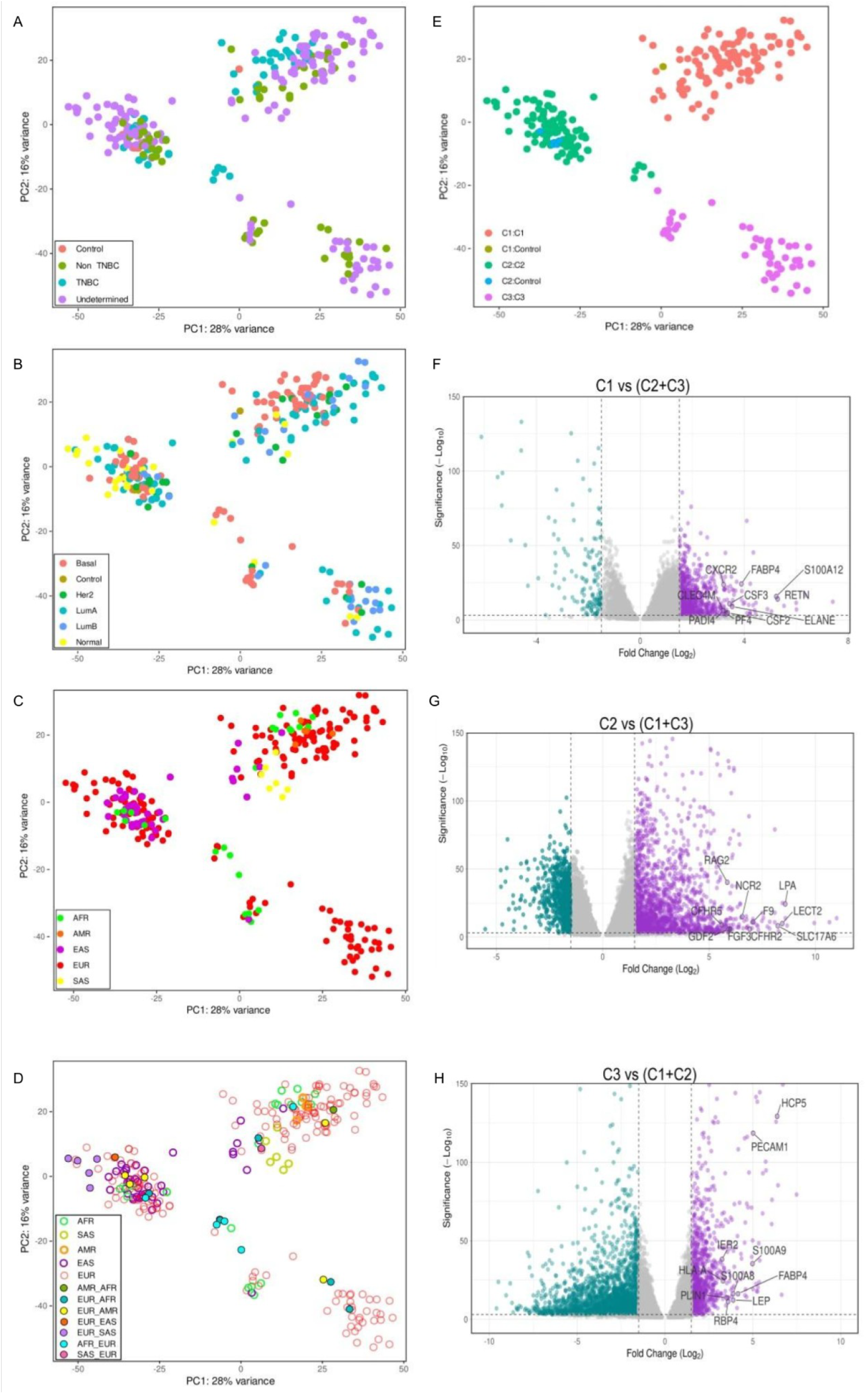
(A-E) PCA plot of gene counts with VST for 280 RNA-seq samples using 49,059 genes. A. Colored by TNBC status. B. Colored by PAM50 subtypes. C. Colored by five major human population predictions. (D) Colored by human population filling dots for admixed individuals. (E) colored by K-means clustering. Each dot represents one sample. (F-H) Volcano plots of differentially expressed genes (DEGs) for the cluster comparisons (F) C1 vs (C2+C3) (G), C2 vs (C1+C3) (H), and C3 vs (C1+C2). Upregulated genes were highlighted in purple and downregulated genes in teal. Key immune-related genes were annotated based on their known functional relevance, to emphasize their biological importance in each comparison.

Figure 4E shows the k-means clustering of the samples if three groups are set as a parameter. To investigate the molecular distinctions among the newly defined clusters (termed C1, C2 and C3), differential expression analysis was performed on the three clusters, comparing each cluster with the other two (set as controls). Figures 4F-4H show the Volcano plots corresponding to the three comparisons, i.e. C1 vs (C2+C3), C2 vs (C1+C3), and C3 vs (C1+C2). The plots highlight key upregulated genes that significantly influence immune regulation, inflammation, and metabolic processes. In the C1 vs (C2+C3) comparison (Figure 4F), genes associated with immune activation were prominently upregulated, including S100A12, CXCR2, CSF3, and RETN. These genes are involved in pathways such as cytokine signaling, natural killer cell activation, and inflammatory responses. Notably, FABP4 and PADI4 were also upregulated, reflecting a link between lipid metabolism and immune processes. These findings suggest that Cluster 1 is characterized by strong immune activation and inflammatory responses. For the C2 vs (C1+C3) comparison (Figure 4G), immune-related genes such as RAG2, NCR2, and CFHR5 were among the most significantly upregulated. These genes are known to play key roles in lymphocyte development, complement regulation, and immune signaling. Additionally, genes like LPA and FGF3, which are involved in lipid metabolism and cellular signaling, further indicate the involvement of metabolic-immune crosstalk in Cluster 2. In the C3 vs (C1+C2) comparison (Figure 4H), genes such as HCP5, PECAM1, HLA-A, S100A8, and LEP were significantly upregulated. These genes are linked to antigen presentation, immune modulation, and inflammation, processes that suggest an immunologically active and metabolically regulated environment in Cluster 3. FABP4 and S100A9, known regulators of inflammation and immune responses, further support the distinctive immune landscape of this cluster.

We performed a functional enrichment analysis comparing Cluster 1 against Clusters 2 and 3 to investigate the functional implications of gene expression differences among clusters. The results are summarized in Figure 5. In the C1 vs (C2+C3) comparison (Figure 5A), pathways associated with metabolic processes were prominently enriched, including “Drug metabolism by cytochrome P450” and “Retinol metabolism,” alongside immune-related processes such as “Natural killer cell-mediated cytotoxicity” and “JAK-STAT signaling pathway.” This suggests a strong involvement of metabolic reprogramming and immune activation in Cluster C1. For the C2 vs (C1+C3) comparison (Figure 5B), pathways related to immune signaling were significantly enriched, including the “Cytosolic DNA-sensing pathway,” “RIG-I-like receptor signaling pathway,” and “Cytokine-cytokine receptor interaction,” highlighting a robust immune response in Cluster C2. Additionally, pathways such as “Chemical carcinogenesis-DNA adducts” and “Drug metabolism-cytochrome P450” reflect metabolic and chemical stress adaptations. Lastly, the C3 vs (C1+C2) comparison (Figure 5C) revealed significant enrichment for pathways involved in immune regulation and inflammation, such as “Allograft rejection,” “IL-17 signaling pathway,” and “Antigen processing and presentation.” Notably, processes related to cellular senescence were also enriched, indicating a potential role for immune suppression and aging-related mechanisms in Cluster C3.

**Figure 5.**
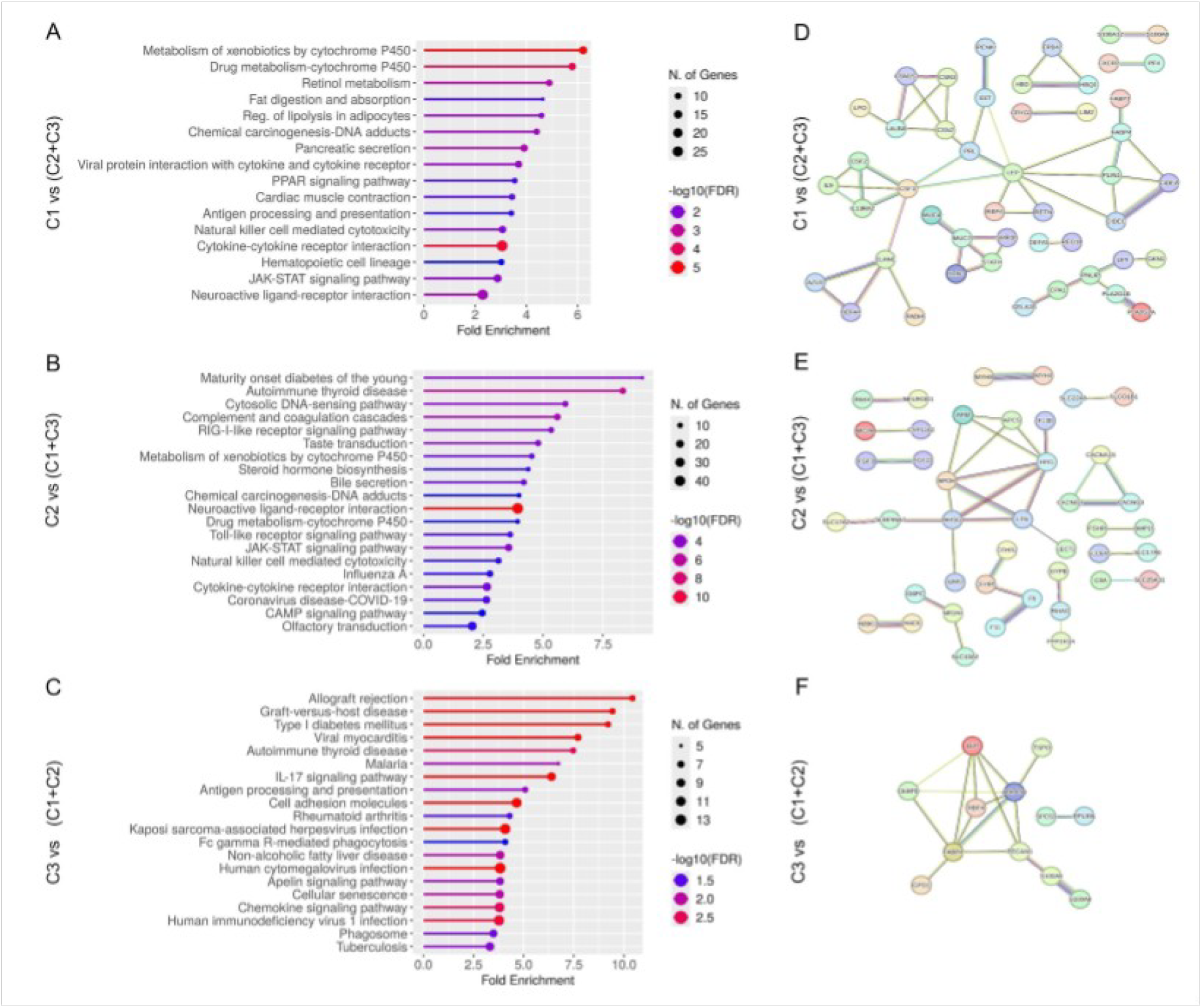
A, B & C: Functional enrichment analysis of upregulated DEGs for cluster comparisons. Bar plots illustrate the enriched KEGG pathways for the comparisons C1 vs (C2+C3) (A), C2 vs (C1+C3) (B), and C3 vs (C1+C2) (C). The x-axis represents fold enrichment, indicating the magnitude of pathway enrichment relative to the background gene set. The size of the dots corresponds to the number of genes contributing to each pathway, while the color gradient reflects statistical significance as -log10(FDR), with darker shades representing more significant results. This visualization highlights distinct pathway enrichments across the comparisons, emphasizing processes such as metabolism, immune signaling, and cellular regulation. Protein-protein interaction (PPI) networks for the top-upregulated DEGs in cluster comparisons. D, E & F: PPI networks were generated using STRING for the comparisons C1 vs (C2+C3) (D), C2 vs (C1+C3) (E), and C3 vs (C1+C2) (F) with a high confidence interaction score of 0.7. Nodes represent upregulated proteins, while edges indicate predicted interactions. Key hub proteins include LEP and CSF2 in C1 vs (C2+C3), APOH and LPA in C2 vs (C1+C3), and LEP, RBP4, and ADIPOQ in C3 vs (C1+C2), highlighting immune, lipid metabolism, and adipokine signaling processes as central to cluster-specific molecular interactions.

To explore the functional relationships among the top upregulated differentially expressed genes (DEGs) for the newly grouped clusters, we performed a protein-protein interaction (PPI) analysis using STRING. The resulting networks revealed distinct interaction patterns across comparisons (Figure 5D-5F). In the C1 vs (C2+C3) comparison (Figure 5D), a highly interconnected network was observed, with key hub proteins such as LEP (leptin) and CSF2 (colony-stimulating factor 2), which are linked to immune regulation and inflammatory processes. For C2 vs (C1+C3) (Figure 5E), hub nodes such as APOH (apolipoprotein H) and LPA (lipoprotein (A)) emerged, highlighting pathways associated with lipid metabolism and coagulation cascades. In contrast, the C3 vs (C1+C2) comparison (Figure 5F) showed a more compact network centered around LEP, RBP4 (retinol-binding protein 4), and ADIPOQ (adiponectin), proteins strongly associated with metabolic regulation and adipokine signaling. These results underscore the cluster-specific molecular interactions, linking immune response and metabolic pathways to the functional differences observed in the enriched gene sets.

### Immune cell composition and *in silico* estimation of drug sensitivities in clustered transcriptomic profiles

To explore potential differences in immune cell infiltration across the three identified clusters, we employed the xCell deconvolution method (Aran et al., 2017), which estimates the frequencies of 64 unique immune and stromal cell populations from bulk transcriptomic profiles. Among these, six immune cell populations displayed significant differences between clusters, highlighting distinct immune microenvironment patterns (Figures 6A to 6F). The cluster C2 exhibited significantly higher infiltration of natural killer (NK) cells, plasma cells, T-helper 1 (Th1) cells, and CD8+ effector memory T (Tem) cells compared to the clusters C1 and C3. These cell types are known for their critical roles in antitumor immunity. For example, NK cells mediate direct cytotoxicity against tumor cells, Th1 cells secrete cytokines like interferon-gamma (IFN-γ) to activate macrophages and enhance the cytotoxic functions of T cells, while CD8+ Tem cells are vital for sustained antitumor immune responses. The elevated infiltration of these cells in C2 suggests that this cluster may represent a more immunoreactive phenotype, potentially more responsive to immunotherapies.

**Figure 6.**
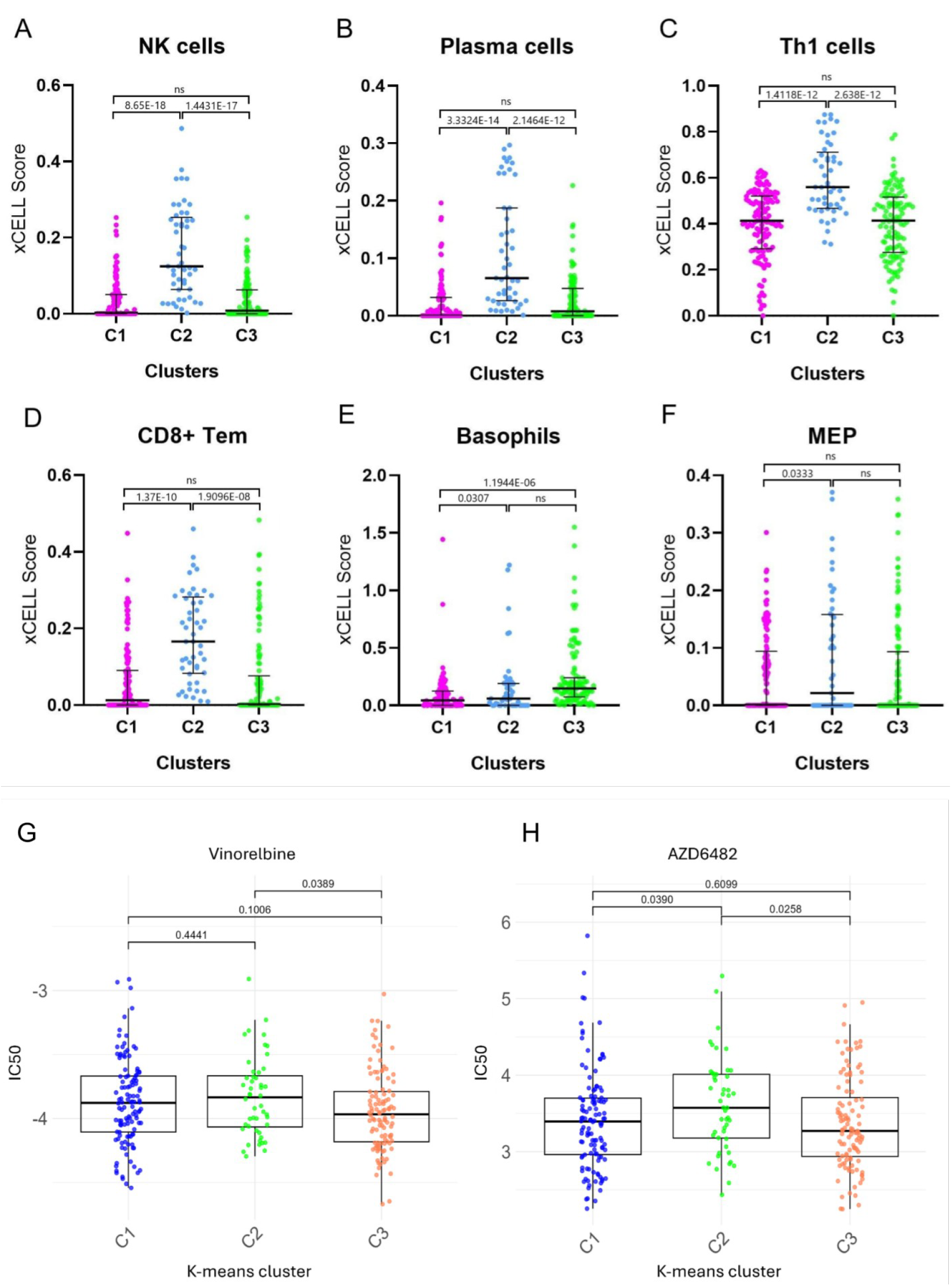
(A-F) Differences in immune cell infiltration among the three identified clusters (C1, C2, and C3) based on xCell deconvolution analysis. The xCell scores represent the estimated frequencies of six immune cell populations: NK cells (Natural Killer cells), Plasma cells, Th1 cells (T-helper 1 cells), CD8+ Tem cells (CD8+ Effector Memory T cells), Basophils, and MEP cells (Megakaryocyte-Erythroid Progenitors). Paired comparisons between clusters were performed using the Mann-Whitney U test, with p-values < 0.05 considered significant. Error bars represent the median and interquartile range. (G,H) Drug sensitivity predictions for FDA-approved drugs across transcriptomic clusters using the pRRophetic R package. (G) Distribution of predicted IC50 values for Vinorelbine, an antimitotic agent. (H) Distribution of predicted IC50 values for AZD6482, a PI3Kβ inhibitor. Statistical significance was determined using pairwise comparisons, and p-values are indicated above the boxplots.

Additionally, basophils were significantly elevated in C1, compared to C2 and C3, suggesting a distinct immune profile. While basophils are typically associated with allergic responses, their role in cancer is context-dependent, as they can either promote or inhibit tumor progression depending on the microenvironment. Lastly, megakaryocyte-erythroid progenitor (MEP) cells showed moderate differences, with slightly higher levels in C2 compared to other clusters. The role of these cells in shaping the immune landscape in cancer is currently unknown.

Breast cancer remains a highly heterogeneous disease, with varying responses to standardized treatments across molecular subtypes. While chemotherapy is effective for many patients, challenges persist in cases of refractory disease and particularly in triple-negative breast cancer (TNBC), where therapeutic options remain limited, and there is a critical need for novel treatments. In this study, we addressed this need by exploring potential drug sensitivities through an *in silico* prediction of response to FDA-approved drugs based on transcriptomic profiles from individual patient clusters. The analysis revealed significant differences in predicted drug sensitivities across the clusters. Specifically, the cluster C3 showed higher sensitivity (lower IC50) to Vinorelbine, compared to the cluster C2 (Figure 6G). Vinorelbine is an antimitotic agent that disrupts microtubule dynamics to induce mitotic arrest and cell death. It has been widely used for treatment of breast cancer and other malignancies. C2 demonstrated lower sensitivity to AZD6482, compared to C1 and C3 (Figure 6H). AZD6482 is a PI3Kβ inhibitor that targets key signaling pathways critical for cell growth and survival. It has potential activity in tumors driven by PI3K signaling dysregulation, including TNBC (p = 0.0258).

## Discussion

Given the worldwide importance of breast cancer (BC) as a public health problem, the biology of BC tumors and the development of methodologies for differential identification and diagnosis have been active research topics for more than 25 years (Perou et al., 2000; Lawton et al., 2023). In this work, we aimed to provide new knowledge on functional genomics and classification of BC tumors by bioinformatic reanalysis of 274 RNA-seq tumors and six healthy tissue samples obtained from public databases, originally published by a wide range of studies sampling different populations and having different research goals. The use of current tools for genotyping, machine learning, differential expression, and cell type deconvolution revealed novel information on the estimation of genetic ancestries, clustering and genes with differential expression among patients, leading to predictions of differential treatments.

Molecular subtyping of these samples following the PAM50 method displayed ambiguous molecular profiles with high heterogeneity. A large number of samples were assigned to more than one group. One of the limitations of the PAM50 model is that the intrinsic subtypes are not discrete categories, but they represent a continuous of molecular characteristics. For example, Luminal A can share molecular characteristics with Normal-like, which translates into a membership probability of belonging to both subtypes for samples belonging to either subtype. These findings align with recent studies highlighting the limitations of PAM50 in accurately capturing the molecular diversity of breast cancer, particularly in non-European populations (Okimoto et al., 2024). This issue is critical because inaccurate subtype classification can lead to inappropriate treatment regimes, such as prescribing ineffective therapies or overlooking targeted options, ultimately worsening patient outcomes. The unsupervised clustering of the samples based on full transcriptome levels did not reveal groups consistent with the current subtypes. This suggests that new classification methods could be developed based on complete expression data and a more inclusive sampling, especially regarding ethnicity. The observed differences between clusters after deconvolution into cell subtypes also suggest that single-cell RNA-seq data can be a great resource to redefine molecular subtypes in breast cancer.

Although RNA-seq has been available for more than ten years as a tool to obtain a complete characterization of gene expression profiles, clinical diagnosis still relies on data obtained from chip-based methods targeting a limited number of genes. This is understandable because cost-benefit is an important aspect to consider when making decisions on methods for diagnosis. However, the reductions in sequencing costs and the amount of information that can be obtained in RNA-seq experiments suggest that RNA-seq can become a direct BC subtyping procedure in the near future. Our results show that broad genetic ancestry can be inferred directly by genotyping selected SNP markers from the aligned RNA-seq reads. This feature enables researchers to obtain genomic variation information in contexts in which it is not feasible or cost-effective to generate DNA sequencing data (Barral-Arca et al., 2019, Razi et al., 2024, Fachrul et al., 2023, Yépez et al., 2022). Moreover, a definition of new subtypes without biases towards particular genes and the analysis of complete transcriptome profiles can be envisioned as a significant step forward toward accurate diagnosis and treatment of cancer patients.

The complete dataset included samples from the five major ancestries described in human population genetic studies. However, the representation disparity among ethnicities was evident in this survey. More than 70% of the samples had European ancestry. This disparity is consistent with those observed in genome-wide association studies in which European ancestry is the most prevalent in public databases (Fitipaldi et al., 2023; Troubat et. al., 2024). Although current large sequencing efforts such as the All of Us sequencing program aim to reduce these disparities for genomic data (The All of Us Research Program Genomics Investigators, 2024), similar efforts are needed to obtain diverse expression data for cancer samples.

Unsupervised machine learning analysis of the transcriptome profiles revealed at least three clearly differentiated clusters of tumors. A functional or demographic interpretation of these clusters was challenging because they did not seem to correspond to cancer subtypes or ethnicities. Regarding subtypes, a possible relationship between clusters and the current subtypes is obscured by the observed issues of accuracy on subtype assignments following the PAM50 protocol. Regarding ethnicity, the eight SAS samples included in this study were grouped in C2 being the only ancestry group distributed in only one cluster. The other populations were spread in two or even the three clusters. These findings highlight the substantial heterogeneity within breast cancer. Different analyses were performed aiming to provide functional annotation of each cluster, to predict cell type compositions, and to assess if differential treatments could be predicted for patients among clusters. The clusters showed diverse immune landscapes and varying sensitivities to specific drugs.

The cluster C1 is characterized by a robust immune activation and inflammatory response profile, supported by the upregulation of key genes such as CSF2 and LEP. The enrichment of pathways like JAK-STAT signaling and Natural killer cell-mediated cytotoxicity highlights the involvement of immune regulatory mechanisms that contribute to tumor progression and immune evasion. CSF2 has been implicated in enhancing tumor metastasis via the FAK pathway, and its elevated expression aligns with an aggressive phenotype (He et al., 2023). Similarly, LEP, an adipokine linked to obesity-related cancers, supports angiogenesis and tumor growth (Andò et al., 2012). These findings suggest that C1 tumors are driven by inflammatory and immune dysregulation, which may make them responsive to therapies targeting inflammatory mediators or metabolic pathways.

The cluster C2 displays a distinct metabolic profile with adaptive metabolic responses to chemical stress, reflected by the upregulation of genes involved in Cytochrome P450 drug metabolism and Steroid Hormone Biosynthesis pathways. These pathways are known to influence carcinogenesis and chemoresistance (Luo et al., 2021; Valko-Rokytovská et al., 2021). The identification of APOH and LPA as key hub proteins further emphasizes the role of lipid metabolism and coagulation in fostering a pro-tumorigenic environment. Intriguingly, C2 also exhibited elevated levels of immune cell infiltration, including NK cells, Th1 cells, and CD8+ Tem cells, suggesting a more immunoreactive phenotype. The presence of these immune cell types, known for their anticancer roles, indicates that tumors in this cluster may be more responsive to immunotherapy. Future studies can be directed to explore the potential for combining immunotherapy with metabolic pathway inhibitors in this cluster.

Tumors within the cluster C3 exhibit a unique combination of inflammatory and metabolic processes, with significant enrichment of pathways like IL-17 signaling” and “Chemokine signaling. These pathways contribute to the recruitment of immune and stromal cells, creating an inflammatory tumor microenvironment that promotes immune evasion and metastasis (Amatya et al., 2017; Hembruff et al., 2009). LEP and RBP4 were selected as central hub proteins, highlighting the importance of adipokine signaling in this cluster. C3 also showed higher sensitivity to vinorelbine, an antimitotic agent, consistent with its known efficacy in breast cancer treatment (Aapro et al., 2012). The higher sensitivity of C3 to AZD6482, a PI3Kβ inhibitor, aligns with the dysregulation of PI3K signaling frequently observed in breast cancer (Miricescu et al., 2020). These findings suggest that C3 tumors may benefit from targeted therapies addressing inflammatory and metabolic dysregulation, alongside drug repurposing efforts.

The recent development of single cell RNA sequencing (scRNA-seq) provides a new level of detail to investigate expression profiles of breast cancer and other tumors. Although the cost of this technique is prohibitively large for direct clinical use, it is possible to perform deconvolution of bulk expression profiles taking advantage of publicly available cell type specific profiles obtained from scRNA-seq experiments. This allowed us to extract novel information of publicly available bulk RNA-seq data regarding immunological heterogeneity among clusters. In particular, the immunoreactive phenotype of C2 exhibited significantly higher infiltration of NK cells, Th1 cells, and CD8+ Tem cells, compared to other clusters. These immune cell types have been associated with improved patient outcomes and enhanced responses to immunotherapy in breast cancer (Rezaeifard et al., 2021; Nersesian et al., 2021). By integrating immune infiltration data with drug sensitivity predictions, we provide a framework for generating new preclinical therapeutic hypotheses. For instance, the immunoreactive nature of C2 suggests that tumors within this cluster may respond well to immune checkpoint inhibitors. In contrast, tumors within the C3 cluster may benefit from metabolic pathway inhibitors combined with agents such as vinorelbine or AZD6482.

In conclusion, this study demonstrated that genetic ancestry can be predicted with tumor RNA-seq data and the PAM50 subtypes are not consistent with gene expression in the RNA-seq experiment in our samples. Unsupervised analysis of the currently available expression data grouped tumors in clusters without evident relationship with demographic metadata or current subtyping. Functional enrichment, immune profiling and drug sensitivity analysis revealed insights for new tailored therapeutic approaches. These insights provide a foundation for generating hypotheses about personalized treatment strategies for patients with limited therapeutic options. Validation of these insights require the analysis of larger datasets with better balance regarding ethnicity, experimental validation, and clinical studies, in order to translate them into actionable treatment strategies.

## Methods

### Data collection and initial processing

The Human reference genome hg38 was downloaded from the Broad Institute public database (https://console.cloud.google.com/storage/browser/gcp-public-data--broad-references;tab=objects?prefix=&forceOnObjectsSortingFiltering=false). The complete dataset analyzed in this study consists on publicly available RNA-seq data obtained from breast cancer samples of anonymized patients. The data was recovered from the Sequence Read Archive (SRA) database, using the search term “(breast cancer) AND “Homo sapiens”[orgn:txid9606] NOT cell”, then adding as a filter “fastq” in the File type section, “RNA” in Source section and “Public” in Access section. The results were filtered by a minimum number of 10 million reads. We selected a total of 274 samples based on percentage of mapped reads and the geographical origin of the study. We also included 6 samples of healthy tissue that were used as controls, for a dataset of 280 samples in total. RNA-seq reads were mapped to the human reference genome using HISAT2 (v.2.2.1) (Kim, *et al*., 2019) obtaining aligned reads in SAM format. SAM files were sorted using Picard (v2.27.4).

### Ancestry prediction

Regions covered by RNA-seq reads were calculated from BAM files using samtools depth (v.1.16.1) (Li *et al*., 2009) and were joined to generate the set of common regions between all the samples. Genomic variation data from the human reference populations was retrieved from the 1,000 Genomes Project (The 1000 Genomes Project Consortium, 2015) (http://ftp.1000genomes.ebi.ac.uk/vol1/ftp/data_collections/1000G_2504_high_coverage/working/20201028_3202_raw_GT_with_annot), VCF file was filtered retaining the variants covered by RNA-seq common regions, only women individuals, a minimum MAF value of 0.01, a minimum number of genotyped samples of 1,850 using VCFFilter functionality of NGSEP software (Tello et al., 2019), reference and breast cancer samples variants were merged into a single VCF.

The VCF file was converted to plink format using vcftools (v.0.1.16) (Danecek, *et al*., 2011), ADMIXTURE input was generated using plink (v1.9) (Chang, *et al*., 2015). ADMIXTURE (v.1.3.0) (Alexander, *et al*., 2009) analysis was performed for k values between 1 and 10 with a coefficient of variation (CV) value of 20. We used k=5 to determine the population based on the major groups sampled in the 1,000 Genomes Project, allowing the model to predict the native american component in latin american admixed individuals. An additional classification of admixed individuals was made based on membership probability lower than 0.7 to a single population.

### Extraction of read counts from RNA-seq data

Gene counts were assigned with Stringtie (v.1.3.5) (Pertea *et al*., 2015) from mapped sequences, and the obtained matrix was normalized using VST method of DESeq2 R package (Love *et al*., 2014). An R script was built to evaluate batch effects in the counts using principal component analysis, and also to calculate the correlation between average and variance.

### Molecular subtyping prediction

Molecular subtypes were predicted from the normalized gene count matrix using the R package geneFu (v.2.28.0) (Gendoo *et al*., 2016) using a single sample predictor algorithm (PAM50) (https://www.bioconductor.org/packages/release/bioc/vignettes/genefu/inst/doc/genefu.html) (Tibshirani *et al*., 2002).

### Unsupervised machine learning of gene counts

The gene count matrix was manually curated to remove antisense, guide RNA, lncRNA, miRNA, pseudogenes, rRNA, sequence feature, snoRNA, tRNA, and uncharacterized genes based on gff3 annotation file, additionally, only genes with sums in all samples at least 30 were kept. Resulting in 43,548 transcripts/features included in a Principal Component Analysis (PCA) where an unsupervised machine learning model (K-means) was applied using k values of 1 to 4, which was analyzed manually in 2D and 3D to select the most accurate k value.

### Differential expression analysis

Differential expression analysis was performed to identify significantly upregulated and downregulated genes across the cluster comparisons: C1 vs (C2+C3), C2 vs (C1+C3), and C3 vs (C1+C2). Gene expression data was analyzed using the DESeq2 package in R, which estimates log2 fold changes and calculates adjusted p-values (Benjamini-Hochberg correction) to control for false discovery rates (FDR). Genes with an adjusted p-value < 0.05 and an absolute log2 fold change ≥ 2 were considered significantly differentially expressed. Volcano plots and PCA were generated using the ggplot2 package in R to visualize the results (Wickham et al., 2016).

### Functional Enrichment Analysis

Functional enrichment analysis was conducted to identify biological processes and pathways associated with the upregulated differentially expressed genes (DEGs) across the cluster comparisons: C1 vs (C2+C3), C2 vs (C1+C3), and C3 vs (C1+C2). KEGG pathway enrichment was performed using the clusterProfiler package in R, enabling systematic analysis and visualization of enriched pathways. To ensure precision and biological relevance, the analysis focused on upregulated DEGs with a log fold change of ≥ 2. Enrichment results were evaluated using fold enrichment and adjusted p-values (-log10[FDR]) to determine statistical significance. Data visualization, including enrichment plots, was generated using the ggplot2 package in R (Wickham et al., 2016).

### Protein-Protein Interaction (PPI) Analysis

To identify functional relationships among the upregulated differentially expressed genes (DEGs) in the cluster comparisons, protein-protein interaction (PPI) networks were constructed using the STRING v11.5 database. For each comparison (C1 vs (C2+C3), C2 vs (C1+C3), and C3 vs (C1+C2)), the top 100 significant upregulated DEGs (based on log fold change and adjusted p-value) were selected for analysis. Interactions were filtered using a high-confidence interaction score of 0.7 to ensure reliable associations. The confidence score integrates evidence from experimental data, curated databases, co-expression, and text mining. STRING-generated networks included nodes representing proteins and edges denoting predicted interactions. Edges were color-coded based on the type of evidence supporting the interaction. Networks were visualized to highlight the degree of connectivity, with hub proteins identified as those exhibiting multiple interactions within the network.

### *In Silico* Drug Sensitivity Analysis

Drug sensitivity analysis was performed using the R package pRRophetic (Geeleher et al., 2014), which predicts the half-maximal inhibitory concentration (IC50) of chemotherapeutic agents based on RNA-seq expression data. Sensitivity differences between the K-means clusters (C1, C2, C3) were assessed using the Kruskal-Wallis test, followed by pairwise Wilcoxon rank-sum tests, with p < 0.05 considered significant.

### Immune Infiltration Analysis

Analysis of the Immune Microenvironment Infiltration was conducted to evaluate immune cell infiltration across clusters using the R package **xCell** via its web server (https://xcell.ucsf.edu/). xCell is a computational method that estimates the relative abundance of 64 distinct cell populations, including immune and stromal cell types, based on curated gene expression signatures. Differential infiltration of these populations was analyzed across the clusters, with a significance threshold set at p < 0.05.

## Acknowledgements and Funding

The experiments presented in this manuscript were supported by internal funds from Universidad de los Andes through the masters program in computational biology. We acknowledge the support of the IT Services Department and ExaCore-IT core-facility of the Vice Presidency for Research & Creation at Universidad de Los Andes for their technical support to perform the bioinformatic analysis.

## Author contributions

JS and JD conceived the study. JS built genomic and gene count databases from the RNA-seq data. All authors performed bioinformatic analysis and wrote the manuscript. All authors approved the final version of the manuscript.

### Conflict of Interests

None declared

### Data availability

All data analyzed in this study is available at the Sequence Read Archive (SRA) database. Accession numbers for each sample are available at the supplementary table 1.

## Notes

### Competing Interest Statement

The authors have declared no competing interest.

## References

Aapro M, Finek J. Oral vinorelbine in metastatic breast cancer: a review of current clinical trial results. Cancer Treat Rev. 2012 Apr;38(2):120–6. doi: 10.1016/j.ctrv.2011.05.005. Epub 2011 Jul 13. PMID: 21742438.

Ades F, Zardavas D, Bozovic-Spasojevic I, et al. Luminal B breast cancer: molecular characterization, clinical management, and future perspectives. J Clin Oncol. 2014 Sep 1;32(25):2794–803. doi: 10.1200/JCO.2013.54.1870. Epub 2014 Jul 21. PMID: 25049332.

Amatya N, Garg AV, Gaffen SL. IL-17 Signaling: The Yin and the Yang. Trends Immunol. 2017 May;38(5):310–322. doi: 10.1016/j.it.2017.01.006. Epub 2017 Feb 20. PMID: 28254169; PMCID: PMC5411326.

Andò S, Catalano S. The multifactorial role of leptin in driving the breast cancer microenvironment. Nat Rev Endocrinol. 2011 Nov 15;8(5):263–75. doi: 10.1038/nrendo.2011.184. PMID: 22083089.

Aran D, Hu Z, Butte AJ. xCell: digitally portraying the tissue cellular heterogeneity landscape. Genome Biol. 2017 Nov 15;18(1):220. doi: 10.1186/s13059-017-1349-1. PMID: 29141660; PMCID: PMC5688663.

Arpino G, Generali D, Sapino A, et al. Gene expression profiling in breast cancer: a clinical perspective. Breast. 2013 Apr;22(2):109–120. doi: 10.1016/j.breast.2013.01.016. Epub 2013 Feb 23. Erratum in: Breast. 2016 Feb;25:86. Del Matro, Lucia [corrected to Del Mastro, Lucia]. PMID: 23462680.

Barral-Arca R, Pardo-Seco J, Bello X, Martinón-Torres F, Salas A. Ancestry patterns inferred from massive RNA-seq data. RNA. 2019 Jul;25(7):857–868. doi: 10.1261/rna.070052.118. Epub 2019 Apr 22. PMID: 31010885; PMCID: PMC6573782.

Bertucci F, Finetti P, Birnbaum D. Basal breast cancer: a complex and deadly molecular subtype. Curr Mol Med. 2012 Jan;12(1):96–110. doi: 10.2174/156652412798376134. PMID: 22082486; PMCID: PMC3343384.

Cadoo KA, Traina TA, King TA. Advances in molecular and clinical subtyping of breast cancer and their implications for therapy. Surg Oncol Clin N Am. 2013 Oct;22(4):823–40. doi: 10.1016/j.soc.2013.06.006. Epub 2013 Jul 27. PMID: 24012401.

Chang CC, Chow CC, Tellier LC, Vattikuti S, Purcell SM, Lee JJ. Second-generation PLINK: rising to the challenge of larger and richer datasets. Gigascience. 2015 Feb 25;4:7. doi: 10.1186/s13742-015-0047-8. PMID: 25722852; PMCID: PMC4342193.

Choi J, Jung WH, Koo JS. Clinicopathologic features of molecular subtypes of triple negative breast cancer based on immunohistochemical markers. Histol Histopathol. 2012 Nov;27(11):1481–93. doi: 10.14670/HH-27.1481. PMID: 23018247.

Ciriello G, Sinha R, Hoadley KA, Jacobsen AS, Reva B, Perou CM, Sander C, Schultz N. The molecular diversity of Luminal A breast tumors. Breast Cancer Res Treat. 2013 Oct;141(3):409–20. doi: 10.1007/s10549-013-2699-3. Epub 2013 Oct 6. PMID: 24096568; PMCID: PMC3824397.

Dai X, Li T, Bai Z, Yang Y, Liu X, Zhan J, Shi B. Breast cancer intrinsic subtype classification, clinical use and future trends. Am J Cancer Res. 2015 Sep 15;5(10):2929-43. PMID: 26693050; PMCID: PMC4656721.

Danecek P, Auton A, Abecasis G, et al. The variant call format and VCFtools. Bioinformatics. 2011 Aug 1;27(15):2156–8. doi: 10.1093/bioinformatics/btr330. Epub 2011 Jun 7. PMID: 21653522; PMCID: PMC3137218.

Eroles P, Bosch A, Pérez-Fidalgo Ja, Lluch A. Molecular biology in breast cancer: intrinsic subtypes and signaling pathways. Cancer Treat Rev. 2012 Oct;38(6):698–707. doi: 10.1016/j.ctrv.2011.11.005. Epub 2011 Dec 16. PMID: 22178455.

Fachrul M, Karkey A, Shakya M, et al. Direct inference and control of genetic population structure from RNA sequencing data. Commun Biol. 2023 Aug 2;6(1):804. doi: 10.1038/s42003-023-05171-9. PMID: 37532769; PMCID: PMC10397182.

Foulkes WD, Stefansson IM, Chappuis PO, Bégin LR, Goffin JR, Wong N, Trudel M, Akslen LA. Germline BRCA1 mutations and a basal epithelial phenotype in breast cancer. J Natl Cancer Inst. 2003 Oct 1;95(19):1482–5. doi: 10.1093/jnci/djg050. PMID: 14519755.

Fejerman L, John EM, Huntsman S, Beckman K, Choudhry S, Perez-Stable E, Burchard EG, Ziv E. Genetic ancestry and risk of breast cancer among U.S. Latinas. Cancer Res. 2008 Dec 1;68(23):9723–8. doi: 10.1158/0008-5472.CAN-08-2039. PMID: 19047150; PMCID: PMC2674787.

Fitipaldi H, Franks PW. Ethnic, gender and other sociodemographic biases in genome-wide association studies for the most burdensome non-communicable diseases: 2005-2022. Hum Mol Genet. 2023 Jan 13;32(3):520–532. doi: 10.1093/hmg/ddac245. PMID: 36190496; PMCID: PMC9851743.

Geeleher P, Cox N, Huang RS. pRRophetic: an R package for prediction of clinical chemotherapeutic response from tumor gene expression levels. PLoS One. 2014 Sep 17;9(9):e107468. doi: 10.1371/journal.pone.0107468. PMID: 25229481; PMCID: PMC4167990.

Harris L, Fritsche H, Mennel R, et al. American Society of Clinical Oncology 2007 update of recommendations for the use of tumor markers in breast cancer. J Clin Oncol. 2007 Nov 20;25(33):5287–312. doi: 10.1200/JCO.2007.14.2364. Epub 2007 Oct 22. PMID: 17954709.

Haque R, Ahmed SA, Inzhakova G, Shi J, Avila C, Polikoff J, Bernstein L, Enger SM, Press MF. Impact of breast cancer subtypes and treatment on survival: an analysis spanning two decades. Cancer Epidemiol Biomarkers Prev. 2012 Oct;21(10):1848–55. doi: 10.1158/1055-9965.EPI-12-0474. Epub 2012 Sep 18. PMID: 22989461; PMCID: PMC3467337.

He X, Wang L, Li H, Liu Y, Tong C, Xie C, Yan X, Luo D, Xiong X. CSF2 upregulates CXCL3 expression in adipocytes to promote metastasis of breast cancer via the FAK signaling pathway. J Mol Cell Biol. 2023 Aug 3;15(4):mjad025. doi: 10.1093/jmcb/mjad025. PMID: 37073091; PMCID: PMC10686244.

Hembruff SL, Cheng N. Chemokine signaling in cancer: Implications on the tumor microenvironment and therapeutic targeting. Cancer Ther. 2009 Apr 14;7(A):254-267. PMID: 20651940; PMCID: PMC2907742.

Howlader N, Altekruse SF, Li CI, Chen VW, Clarke CA, Ries LA, Cronin KA. US incidence of breast cancer subtypes defined by joint hormone receptor and HER2 status. J Natl Cancer Inst. 2014 Apr 28;106(5):dju055. doi: 10.1093/jnci/dju055. PMID: 24777111; PMCID: PMC4580552.

Huo D, Hu H, Rhie SK, et al. Comparison of Breast Cancer Molecular Features and Survival by African and European Ancestry in The Cancer Genome Atlas. JAMA Oncol. 2017 Dec 1;3(12):1654–1662. doi: 10.1001/jamaoncol.2017.0595. PMID: 28472234; PMCID: PMC5671371.

Jack RH, Møller H, Robson T, Davies EA. Breast cancer screening uptake among women from different ethnic groups in London: a population-based cohort study. BMJ Open. 2014 Oct 16;4(10):e005586. doi: 10.1136/bmjopen-2014-005586. PMID: 25324320; PMCID: PMC4202018.

Januszewski A, Tanna N, Stebbing J. Ethnic variation in breast cancer incidence and outcomes--the debate continues. Br J Cancer. 2014 Jan 7;110(1):4–6. doi: 10.1038/bjc.2013.775. PMID: 24398563; PMCID: PMC3887313.

Joyce DP, Murphy D, Lowery AJ, Curran C, Barry K, Malone C, McLaughlin R, Kerin MJ. Prospective comparison of outcome after treatment for triple-negative and non-triple-negative breast cancer. Surgeon. 2017 Oct;15(5):272–277. doi: 10.1016/j.surge.2016.10.001. Epub 2016 Oct 27. PMID: 28277293.

Jung J, Kang E, Gwak JM, Seo AN, Park SY, Lee AS, Baek H, Chae S, Kim EK, Kim SW. Association between basal-like phenotype and BRCA1/2 germline mutations in Korean breast cancer patients. Curr Oncol. 2016 Oct;23(5):298–303. doi: 10.3747/co.23.3054. Epub 2016 Oct 25. PMID: 27803593; PMCID: PMC5081005.

Kim D, Paggi JM, Park C, Bennett C, Salzberg SL. Graph-based genome alignment and genotyping with HISAT2 and HISAT-genotype. Nat Biotechnol. 2019 Aug;37(8):907–915. doi: 10.1038/s41587-019-0201-4. Epub 2019 Aug 2. PMID: 31375807; PMCID: PMC7605509.

Lawton TJ. Update on the Use of Molecular Subtyping in Breast Cancer. Adv Anat Pathol. 2023 Nov 1;30(6):368–373. doi: 10.1097/PAP.0000000000000416. Epub 2023 Sep 25. PMID: 37746905.

Li E, Guida JL, Tian Y, et al. Associations between mammographic density and tumor characteristics in Chinese women with breast cancer. Breast Cancer Res Treat. 2019 Sep;177(2):527–536. doi: 10.1007/s10549-019-05325-6. Epub 2019 Jun 28. PMID: 31254158; PMCID: PMC7304859.

Li H, Handsaker B, Wysoker A, Fennell T, et al. The Sequence Alignment/Map format and SAMtools. Bioinformatics. 2009 Aug 15;25(16):2078–9. doi: 10.1093/bioinformatics/btp352. Epub 2009 Jun 8. PMID: 19505943; PMCID: PMC2723002.

Love MI, Huber W, Anders S. Moderated estimation of fold change and dispersion for RNA-seq data with DESeq2. Genome Biol. 2014;15(12):550. doi: 10.1186/s13059-014-0550-8. PMID: 25516281; PMCID: PMC4302049.

Luo B, Yan D, Yan H, Yuan J. Cytochrome P450: Implications for human breast cancer. Oncol Lett. 2021 Jul;22(1):548. doi: 10.3892/ol.2021.12809. Epub 2021 May 24. PMID: 34093769; PMCID: PMC8170261.

Marker KM, Zavala VA, Vidaurre T, et al. Human Epidermal Growth Factor Receptor 2-Positive Breast Cancer Is Associated with Indigenous American Ancestry in Latin American Women. Cancer Res. 2020 May 1;80(9):1893–1901. doi: 10.1158/0008-5472.CAN-19-3659. Epub 2020 Apr 3. PMID: 32245796; PMCID: PMC7202960.

Mathews JC, Nadeem S, Levine AJ, Pouryahya M, Deasy JO, Tannenbaum A. Robust and interpretable PAM50 reclassification exhibits survival advantage for myoepithelial and immune phenotypes. NPJ Breast Cancer. 2019 Sep 9;5:30. doi: 10.1038/s41523-019-0124-8. PMID: 31531391; PMCID: PMC6733897.

Mavaddat N, Barrowdale D, Andrulis IL, et al. Pathology of breast and ovarian cancers among BRCA1 and BRCA2 mutation carriers: results from the Consortium of Investigators of Modifiers of BRCA1/2 (CIMBA). Cancer Epidemiol Biomarkers Prev. 2012 Jan;21(1):134–47. doi: 10.1158/1055-9965.EPI-11-0775. Epub 2011 Dec 5. PMID: 22144499; PMCID: PMC3272407.

Miricescu D, Totan A, Stanescu-Spinu II, Badoiu SC, Stefani C, Greabu M. PI3K/AKT/mTOR Signaling Pathway in Breast Cancer: From Molecular Landscape to Clinical Aspects. Int J Mol Sci. 2020 Dec 26;22(1):173. doi: 10.3390/ijms22010173. PMID: 33375317; PMCID: PMC7796017.

Nedeljković M, Damjanović A. Mechanisms of Chemotherapy Resistance in Triple-Negative Breast Cancer-How We Can Rise to the Challenge. Cells. 2019 Aug 22;8(9):957. doi: 10.3390/cells8090957. PMID: 31443516; PMCID: PMC6770896.

Nersesian S, Schwartz SL, Grantham SR, MacLean LK, Lee SN, Pugh-Toole M, Boudreau JE. NK cell infiltration is associated with improved overall survival in solid cancers: A systematic review and meta-analysis. Transl Oncol. 2021 Jan;14(1):100930. doi: 10.1016/j.tranon.2020.100930. Epub 2020 Nov 10. PMID: 33186888; PMCID: PMC7670197.

Norum JH, Andersen K, Sørlie T. Lessons learned from the intrinsic subtypes of breast cancer in the quest for precision therapy. Br J Surg. 2014 Jul;101(8):925–38. doi: 10.1002/bjs.9562. Epub 2014 May 21. PMID: 24849143.

Ochoa S, de Anda-Jáuregui G, Hernández-Lemus E. Multi-Omic Regulation of the PAM50 Gene Signature in Breast Cancer Molecular Subtypes. Front Oncol. 2020 May 22;10:845. doi: 10.3389/fonc.2020.00845. PMID: 32528899; PMCID: PMC7259379.

Okimoto LYS, Mendonca-Neto R, Nakamura FG, Nakamura EF, Fenyö D, Silva CT. Few-shot genes selection: subset of PAM50 genes for breast cancer subtypes classification. BMC Bioinformatics. 2024 Mar 1;25(1):92. doi: 10.1186/s12859-024-05715-8. PMID: 38429657; PMCID: PMC10908178.

Parker JS, Mullins M, Cheang MC, et al. Supervised risk predictor of breast cancer based on intrinsic subtypes. J Clin Oncol. 2009 Mar 10;27(8):1160–7. doi: 10.1200/JCO.2008.18.1370. Epub 2009 Feb 9. PMID: 19204204; PMCID: PMC2667820.

Parise CA, Bauer KR, Caggiano V. Variation in breast cancer subtypes with age and race/ethnicity. Crit Rev Oncol Hematol. 2010 Oct;76(1):44–52. doi: 10.1016/j.critrevonc.2009.09.002. Epub 2009 Oct 2. PMID: 19800812.

Perou CM, Sørlie T, Eisen MB, et al. Molecular portraits of human breast tumours. Nature. 2000 Aug 17;406(6797):747–52. doi: 10.1038/35021093. PMID: 10963602.

Pertea M, Pertea GM, Antonescu CM, Chang TC, Mendell JT, Salzberg SL. StringTie enables improved reconstruction of a transcriptome from RNA-seq reads. Nat Biotechnol. 2015 Mar;33(3):290–5. doi: 10.1038/nbt.3122. Epub 2015 Feb 18. PMID: 25690850; PMCID: PMC4643835.

Petros WP, Hopkins PJ, Spruill S, et al. Associations between drug metabolism genotype, chemotherapy pharmacokinetics, and overall survival in patients with breast cancer. J Clin Oncol. 2005 Sep 1;23(25):6117–25. doi: 10.1200/JCO.2005.06.075. Epub 2005 Aug 8. PMID: 16087946.

Prat A, Pineda E, Adamo B, Galván P, Fernández A, Gaba L, Díez M, Viladot M, Arance A, Muñoz M. Clinical implications of the intrinsic molecular subtypes of breast cancer. Breast. 2015 Nov;24 Suppl 2:S26–35. doi: 10.1016/j.breast.2015.07.008. Epub 2015 Aug 5. PMID: 26253814.

Razi A, Lo CC, Wang S, Leek JT, Hansen KD. Genotype prediction of 336,463 samples from public expression data. bioRxiv [Preprint]. 2024 Mar 13:2023.10.21.562237. doi: 10.1101/2023.10.21.562237. PMID: 38559266; PMCID: PMC10979922.

Rezaeifard S, Talei A, Shariat M, Erfani N. Tumor infiltrating NK cell (TINK) subsets and functional molecules in patients with breast cancer. Mol Immunol. 2021 Aug;136:161–167. doi: 10.1016/j.molimm.2021.03.003. Epub 2021 Jun 23. PMID: 34171565.

Roelands J, Mall R, Almeer H, Thomas R, Mohamed MG, Bedri S, Al-Bader SB, Junejo K, Ziv E, Sayaman RW, Kuppen PJK, Bedognetti D, Hendrickx W, Decock J. Ancestry-associated transcriptomic profiles of breast cancer in patients of African, Arab, and European ancestry. NPJ Breast Cancer. 2021 Feb 8;7(1):10. doi: 10.1038/s41523-021-00215-x. PMID: 33558495; PMCID: PMC7870839.

Ross JS, Hatzis C, Symmans WF, Pusztai L, Hortobágyi GN. Commercialized multigene predictors of clinical outcome for breast cancer. Oncologist. 2008 May;13(5):477–93. doi: 10.1634/theoncologist.2007-0248. Erratum in: Oncologist. 2008 Aug;13(8). doi: 10.1634/theoncologist.2007-0248. PMID: 18515733.

Schaafsma E, Zhang B, Schaafsma M, Tong CY, Zhang L, Cheng C. Impact of Oncotype DX testing on ER+ breast cancer treatment and survival in the first decade of use. Breast Cancer Res. 2021 Jul 17;23(1):74. doi: 10.1186/s13058-021-01453-4. PMID: 34274003; PMCID: PMC8285794.

Shah PD, Nathanson KL. Application of Panel-Based Tests for Inherited Risk of Cancer. Annu Rev Genomics Hum Genet. 2017 Aug 31;18:201–227. doi: 10.1146/annurev-genom-091416-035305. Epub 2017 May 15. PMID: 28504904.

Su Y, Zheng Y, Zheng W, Gu K, Chen Z, Li G, Cai Q, Lu W, Shu XO. Distinct distribution and prognostic significance of molecular subtypes of breast cancer in Chinese women: a population-based cohort study. BMC Cancer. 2011 Jul 12;11:292. doi: 10.1186/1471-2407-11-292. PMID: 21749714; PMCID: PMC3157458.

Sung H, Ferlay J, Siegel RL, Laversanne M, Soerjomataram I, Jemal A, Bray F. Global Cancer Statistics 2020: GLOBOCAN Estimates of Incidence and Mortality Worldwide for 36 Cancers in 185 Countries. CA Cancer J Clin. 2021 May;71(3):209–249. doi: 10.3322/caac.21660. Epub 2021 Feb 4. PMID: 33538338.

Szymiczek A, Lone A, Akbari MR. Molecular intrinsic versus clinical subtyping in breast cancer: A comprehensive review. Clin Genet. 2021 May;99(5):613–637. doi: 10.1111/cge.13900. Epub 2020 Dec 28. PMID: 33340095.

Tello D, Gil J, Loaiza CD, Riascos JJ, Cardozo N, Duitama J. NGSEP3: accurate variant calling across species and sequencing protocols. Bioinformatics. 2019 Nov 1;35(22):4716–4723. doi: 10.1093/bioinformatics/btz275. PMID: 31099384; PMCID: PMC6853766.

Tekesin K, Emin Gunes M, Bayrak S, Akar E, Ozturk T, Altinay S, Tural D. PTEN loss is a predictive marker for HER2-positive metastatic breast cancer patients treated with trastuzumab-based therapies. J BUON. 2019 Sep-Oct;24(5):1920-1926. PMID: 31786856.

The All of Us Research Program Genomics Investigators. Genomic data in the All of Us Research Program. Nature 627, 340–346 (2024). 10.1038/s41586-023-06957-x

Telli ML, Chang ET, Kurian AW, et al. Asian ethnicity and breast cancer subtypes: a study from the California Cancer Registry. Breast Cancer Res Treat. 2011 Jun;127(2):471–8. doi: 10.1007/s10549-010-1173-8. Epub 2010 Oct 19. PMID: 20957431; PMCID: PMC4349378.

Tibshirani R, Hastie T, Narasimhan B, Chu G. Diagnosis of multiple cancer types by shrunken centroids of gene expression. Proc Natl Acad Sci U S A. 2002 May 14;99(10):6567–72. doi: 10.1073/pnas.082099299. PMID: 12011421; PMCID: PMC124443.

Troester MA, Sun X, Allott EH, et al. Racial Differences in PAM50 Subtypes in the Carolina Breast Cancer Study. J Natl Cancer Inst. 2018 Feb 1;110(2):176–82. doi: 10.1093/jnci/djx135. PMID: 28859290; PMCID: PMC6059138.

Troubat L, Fettahoglu D, Henches L, Aschard H, Julienne H. Multi-trait GWAS for diverse ancestries: mapping the knowledge gap. BMC Genomics. 2024 Apr 17;25(1):375. doi: 10.1186/s12864-024-10293-3. PMID: 38627641; PMCID: PMC11022331.

Ye J, Xia X, Dong W, Hao H, et al. Cellular uptake mechanism and comparative evaluation of antineoplastic effects of paclitaxel-cholesterol lipid emulsion on triple-negative and non-triple-negative breast cancer cell lines. Int J Nanomedicine. 2016 Aug 24;11:4125–40. doi: 10.2147/IJN.S113638. PMID: 27601899; PMCID: PMC5003597.

Yépez VA, Gusic M, Kopajtich R, et al. Clinical implementation of RNA sequencing for Mendelian disease diagnostics. Genome Med. 2022 Apr 5;14(1):38. doi: 10.1186/s13073-022-01019-9. PMID: 35379322; PMCID: PMC8981716.

Valko-Rokytovská M, Očenáš P, Salayová A, Kostecká Z. Breast Cancer: Targeting of Steroid Hormones in Cancerogenesis and Diagnostics. Int J Mol Sci. 2021 May 30;22(11):5878. doi: 10.3390/ijms22115878. PMID: 34070921; PMCID: PMC8199112.

Wickham H. ggplot2: Elegant Graphics for Data Analysis. Springer-Verlag New York. 2016. ISBN 978-3-319-24277-4, https://ggplot2.tidyverse.org.

Wisniewski JM, Walker B. Association of Simulated Patient Race/Ethnicity With Scheduling of Primary Care Appointments. JAMA Netw Open. 2020 Jan 3;3(1):e1920010. doi: 10.1001/jamanetworkopen.2019.20010. PMID: 31995215; PMCID: PMC6991290.

Zavala VA, Bracci PM, Carethers JM, et al. Cancer health disparities in racial/ethnic minorities in the United States. Br J Cancer. 2021 Jan;124(2):315–332. doi: 10.1038/s41416-020-01038-6. Epub 2020 Sep 9. PMID: 32901135; PMCID: PMC7852513.

